# Localization of chemical synapses and modulatory release sites in the cardiac ganglion of the crab, *Cancer borealis*

**DOI:** 10.1101/2022.04.25.489413

**Authors:** Mara C.P. Rue, Natasha Baas-Thomas, Priya Iyenger, Lara Scaria, Eve Marder

## Abstract

The crustacean cardiac ganglion (CG) comprises nine neurons that provide rhythmic drive to the heart. The CG is the direct target of multiple modulators. Synapsin-like immunoreactivity was found clustered around the somata of the large cells (LC) and in a neuropil at the anterior branch of the CG trunk. This implicates the soma as a key site of synaptic integration, an unusual configuration in invertebrates. Proctolin is an excitatory neuromodulator of the CG and proctolin-like immunoreactivity exhibited partial overlap with putative chemical synapses near the LCs and at the neuropil. A proctolin-like projection was also found in a pair of excitatory nerves entering the CG. GABA-like immunoreactivity was nearly completely colocalized with chemical synapses near the LCs but absent at the anterior branch neuropil. GABA-like projections were found in a pair of inhibitory nerves entering the CG. *Cancer borealis* Allatostatin B1 (CbAST-B1), red pigment concentrating hormone (RPCH) and FLMRFamide-like immunoreactivity each had a unique pattern of staining and co-localization with putative chemical synapses. These results provide morphological evidence that synaptic input is integrated at LC somata in the CG. Our findings provide a topographical organization for some of the multiple inhibitory and excitatory modulators that alter the rhythmic output of this semi-autonomous motor circuit.

## Introduction

All nervous systems rely on a combination of many neurotransmitters and neuromodulators. These signaling molecules change the intrinsic properties of target neurons and/or synapses and thus change the output of the neuronal circuit (Harris-Warrick & Marder, 1991; Marder, 2012; Marder & Bucher, 2007). The location of neuromodulatory input is important for the effect a given modulator will have on a neuronal circuit (Garcia et al., 2015; Magee, 2000; Nusbaum et al., 2017). For instance, the same neuromodulator can have different effects on the same circuit depending on the location and type of release (Stein et al., 2007). Therefore, understanding the structure and localization of different inputs in a neuronal circuit can give insight into how signals are integrated, and how they might affect the neuronal output.

The crustacean cardiac ganglion (CG) generates reliable, stereotyped rhythmic impulses that initiate the contraction of the neurogenic heart, and has been used to study fundamental mechanisms by which rhythmic motor patterns are generated and modulated (Cooke, 2002). Although the CG contains just nine neurons, its stereotyped neuronal output is both flexible and robust (Cooke, 2002; Cruz-Bermúdez & Marder, 2007; Lane et al., 2016; Tazaki & Cooke, 1979). The physiological response of the crustacean cardiac ganglion to various neuromodulators has been well characterized in a variety of species (Cruz-Bermúdez & Marder, 2007; Dickinson et al., 2015; Lane et al., 2018; Yang et al., 2013). In addition to hormonal modulation, the CG is modulated extrinsically by at least three pairs of input nerves (Alexandrowicz, 1934). The anterior pair of nerves inhibits the CG and two more posterior nerves have excitatory effects (Ando & Kuwasawa, 2004; Cooke, 2002; Maynard Jr, 1953; Yazawa & Kuwasawa, 1992).

Early morphological studies of the CG used light microscopy and dye injections to postulate the existence of both chemical and electrical synapses within and between pre-motor and motor neurons of the CG (Alexandrowicz, 1932). The CG neurons are glutamatergic and the small cells project excitatory glutamatergic synapses onto the large motor neurons (Ando & Kuwasawa, 2004; Delgado et al., 2000). Subsequent electron microscopy studies confirmed the existence and organization of chemical and electrical synaptic connections in sections of the CG (Morganelli and Sherman, 1987; Mirolli et al., 1987). Immunohistochemistry studies have visualized some modulatory inputs in whole-mount preparations of CGs in various species. Extensive GABA-like projections are found throughout the CG of the lobster *P. argus* (Delgado et al., 2000; Yang et al., 2013), and nitrous oxide synthase-like immunoreactivity is found in posterior motor neurons and small pacemaker cells of the crab *C. productus* (Scholz et al., 2002). Recent work has used MALDI imaging to characterize the distribution of neuropeptides in the cardiac ganglion of *C. borealis*, and confirmed 316 unique neuropeptides including those studied here (DeLaney & Li, 2020).

Except for the early characterization using electron microscopy, there is little known about the overall structure and localization of chemical synapses within the CG (Morganelli and Sherman, 1987; Mirolli et al., 1987). We therefore stained whole-mount CGs using a monoclonal antibody generated against *Drosophila* synapsins, SYNORF1 antibody, (Klagges et al., 1996). Synapsins are found on the surface of small synaptic vesicles and are highly conserved across species (Evergren et al., 2007; Südhof, 2004). The antibody used in this study has been verified to specifically mark neuropil in confocal studies from *Drosophila* (Klagges et al., 1996) and many crustacean species including *C. borealis* (Bucher et al., 2007; Goeritz et al., 2013; Skiebe & Ganeshina, 2000; Skiebe & Wollenschläger, 2002).

We performed a series of double-immunostains for synapsin and a neuromodulator. There are dozens of neuromodulators known to affect the rhythmic pattern generated by the crustacean CG (Cruz-Bermúdez & Marder, 2007). Here we focused on a subset of both inhibitory and excitatory neuromodulators, all of which have known effects on the output of the CG rhythm *in vitro* and for which we have reliable and verifiable antibodies in the crab *C. borealis:* proctolin, GABA, allatostatin-B1 (CbAST-B1), red pigment concentrating hormone (RPCH) and FLRFamide-like peptide.

## Methods

### Animals and Dissection

Male Jonah crabs, *C. borealis*, were obtained from Commerical Lobster (N=60 crabs, Boston, MA) and housed in artificial seawater tanks at 10-12°C. On average, animals were acclimated at this temperature for one week before use. Prior to dissection, animals were placed on ice for at least 30 minutes. Dissections were performed as previously described (Cruz-Bermúdez & Marder, 2007). In short, the heart was dissected from the animal and the intact cardiac ganglion (CG) was removed, preserving the connections between the small cells and the large cells. The CG was pinned in a Sylgard-coated (Dow Corning) dish in physiological saline.

### Electrophysiology

Extracellular nerve recordings were made by building wells around nerves using a mixture of Vaseline and 10% mineral oil and placing stainless-steel pin electrodes within the wells to monitor spiking activity. Extracellular nerve recordings were amplified using model 3500 extracellular amplifiers (A-M Systems). Data were acquired using a Digidata 1440 digitizer (Molecular Devices, San Jose) and pClamp data acquisition software (Molecular Devices, San Jose, version 10.5). We stimulated nerves exiting the CG while simultaneously recording the rhythm with an extracellular electrode. For each nerve, stainless steel pin electrodes were placed on either side of the nerve and sealed using Vaseline. Stimuli were delivered using a model 3800 stimulator (A-M Systems). Stimuli were 0.5ms pulses in one second trains with a within-stimulus rate of 20 Hz.

### Methylene Blue Staining

To visualize the cardiac ganglion and innervating fibers more clearly for illustration, some hearts were stained with methylene blue. The heart was removed from the animal and the carapace and opened on the ventral wall. The opened heart was soaked in a solution of 0.3% Methylene Blue dissolved in *C. borealis* saline for 20-25 minutes. Stained preps were then washed with excess saline to remove dye, and pictures were taken using a Leica Wild M5 stereo microscope. When necessary, we repeated the methylene blue staining after careful dissection of muscle to expose covered sections of the CG. For translating photos of methylene blue stains to drawings, we carefully measured the distances between anatomical features and made a representative, proportional illustration.

### Antibodies

Commercially available primary antibodies: anti-GABA (1:250; Sigma-Aldrich A2052), anti-SYNORF (1:50 – 1:250; Developmental Studies Hybridoma Bank 3C11)(Klagges et al., 1996). Peptide CbAST-B1 (VPNDWAHFRGSW-NH_2_) was generated at the Biotechnology Center at the University of Wisconsin–Madison, and sent to Lampire Biological Laboratories, Hypersville, PA, to generate polyclonal rabbit antibodies. Details of the antibody verification and production have been previously reported in *C. borealis* (Szabo et al., 2011). Anti-CbAST-B1 antibody was used at a concentration of 1:500. Rabbit anti-proctolin polyclonal antibodies were used at a concentration of 1:1000 (Davis et al., 1989). FLRFamide-like immunoreactivity was examined using either of two rabbit anti-FLRFamide polyclonal antibodies: 671M (Eve Marder et al., 1987) and 231 (O’Donohue et al., 1984) at final dilutions of 1:100 to 1:300. RPCH-like immunoreactivity was labeled with a rabbit polyclonal antibody (gift of R. Elde, University of Minnesota) at a concentration of 1:500 (Fénelon et al., 1999).

### Immunohistochemistry

Samples were fixed with 4% paraformaldehyde in PBS (440mM NaCl, 11mM KCl, 10mM Na_2_H_2_PO_4_, 2mM KH_2_PO_4_) for 40 minutes at room temperature (RT), then washed 3×10 minutes with PBS containing 0.1% Triton (PBS-T). Subsequently, samples were incubated with one or two primary antibodies described above for 1 hour at RT or overnight at 4°C. Following 3×10 minute PBS-T washes, samples were then incubated with secondary antibodies Alexa 488 donkey anti-rabbit (1:1000; Life Technologies A21206) and Alexa 568 donkey anti-mouse (1:1000; Invitrogen A10037) for 1 hour at RT or overnight at 4°C. A final 3×15 minute wash in PBS was performed before mounting the samples in Vectashield (Vector labs).

To ensure the specificity of the above primary antibodies, we performed adsorption controls for anti-GABA, anti-CbAST-B1, anti-proctolin, anti-RPCH and anti-FLRFamide antibodies. The respective primary antibody was first incubated with a saturating concentration of the target molecule (10^−2^ M GABA, 10^−4^ M proctolin, 10^−3^ M CbAST-B1, 10^−4^ M RPCH and 10 ^-5^ M TNRNFLRFamide) for one hour at RT before incubation with the tissue. All other staining techniques were identical to those described above.

### Microscopy and Image Processing

All images were obtained with a Leica SP5 Spectral Confocal Microscope or Zeiss LSM880 Airy Scan Fast Confocal System with 10X or 20X objectives. Multi-field large images were taken with 10% overlap and stitched with the Leica Application Suite or ZEISS ZEN Microscope Software. Z-stacks were obtained using 1 µm or 2 µm step sizes. Images were processed with ImageJ. Figures and anatomical drawings were made using Adobe Illustrator 2022.

## Results

### Anatomy of the Cancer borealis cardiac ganglion

The crustacean heart is neurogenic and is controlled and innervated by the cardiac ganglion, which is found inside the single-chambered heart on the dorsal wall (Fig, 1A). The cardiac ganglion (CG) comprises nine neurons: four small pacemaker interneurons (SC), and five large motor neurons (LC). The location of CG somata as well as the innervation pattern in the heart muscle can vary substantially between species (Cooke, 2002). Given this, we first set out to illustrate accurately the structure of the CG in *Cancer borealis*. The location of all neuronal cell bodies and the major nerves entering and exiting the ganglion as they are embedded in the heart muscle are seen in Figure 1B. We refer to the large motor neurons by their cell number, established by their location within a given CG (LC 1-5). The normal activity pattern of the CG is illustrated in Figure 1C. The small pacemaker neurons initiate bursts of action potentials that drive synchronous bursting of the LCs (Fig. 1C). All neurons in the CG are strongly electrically coupled, and the SCs make excitatory glutamatergic synapses onto the LCs (Fig. 1D).

**Figure 1.**
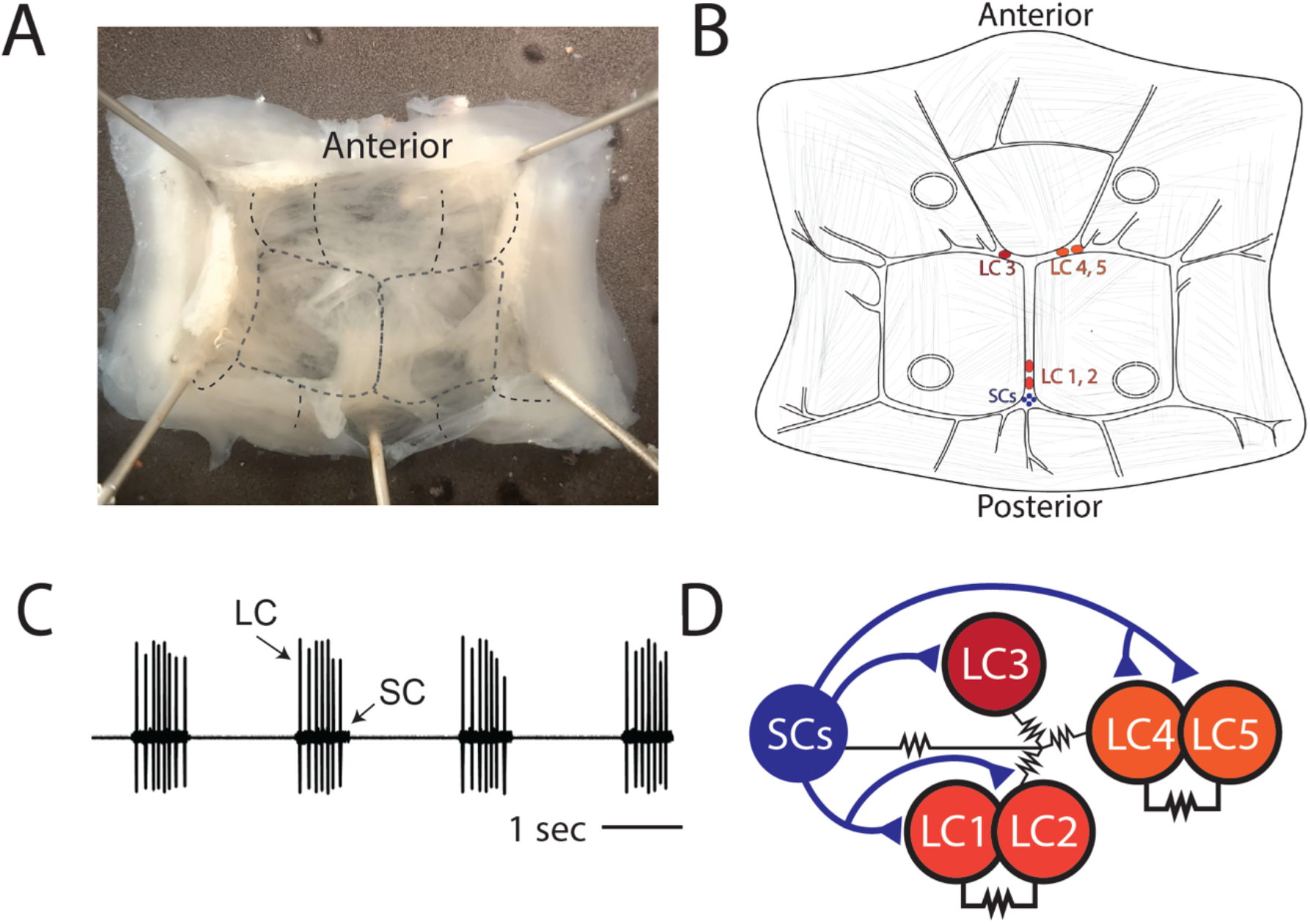
The cardiac ganglion (CG) of crab *Cancer borealis*. A) Crab heart dissected from the animal, opened on the ventral side, and pinned in a dish. The rough location of the CG is outlined in the black dotted line. B) Illustration of the CG showing the locations of the five large motor neurons (LC 1-5, red circles) and the four small interneurons (SC, blue circles). C) Extracellular recording from the trunk of the CG showing the normal synchronous bursting activity of the LC and SC. D) Circuit diagram showing the known chemical and electrical connections between cells in the CG. Triangles indicate excitatory glutamatergic synapses and the resistor symbols indicate electrical coupling.

### Chemical synapses in the CG are clustered around the anterior and posterior branch points and form basket-like structures around large cell bodies

To determine the location and distribution of chemical synapses within the CG, we stained whole-mount CGs using a monoclonal antibody against the pre-synaptic protein, synapsin. Extensive synapsin-like immunoreactivity (synapsin-LIR) is found in several areas of the CG (N=60). Figure 2 illustrates the key aspects of synapsin-LIR in the CG of *C. borealis*, with areas of interest marked in Figure 2A (green boxes). At the anterior Y-shaped branch of the CG there is a dense neuropil of synapsin-LIR that extends along all the branches towards each of the LC soma (Fig. 2B). We also found strikingly dense synapsin-LIR surrounding each of the cell bodies of the anterior large motor neurons (Fig. 2C, D). In addition to small clusters of synapsin-LIR near the LC somata, we found that the synapsin-LIR in basket-like structures that surround and coat the somata. This structure is also found surrounding the posterior LCs (Fig. 2E), where synapsin-LIR surrounds LC1 and LC2. A small neuropil surrounds the small cells, posterior to LC1 and LC2 (Fig. 2E). Apart from the areas shown, we found minimal synapsin-LIR in the rest of the CG. Thus, most putative pre-synaptic terminals within the CG are found along the trunk of the CG and surrounding each of the large motor neuron somata.

**Figure 2.**
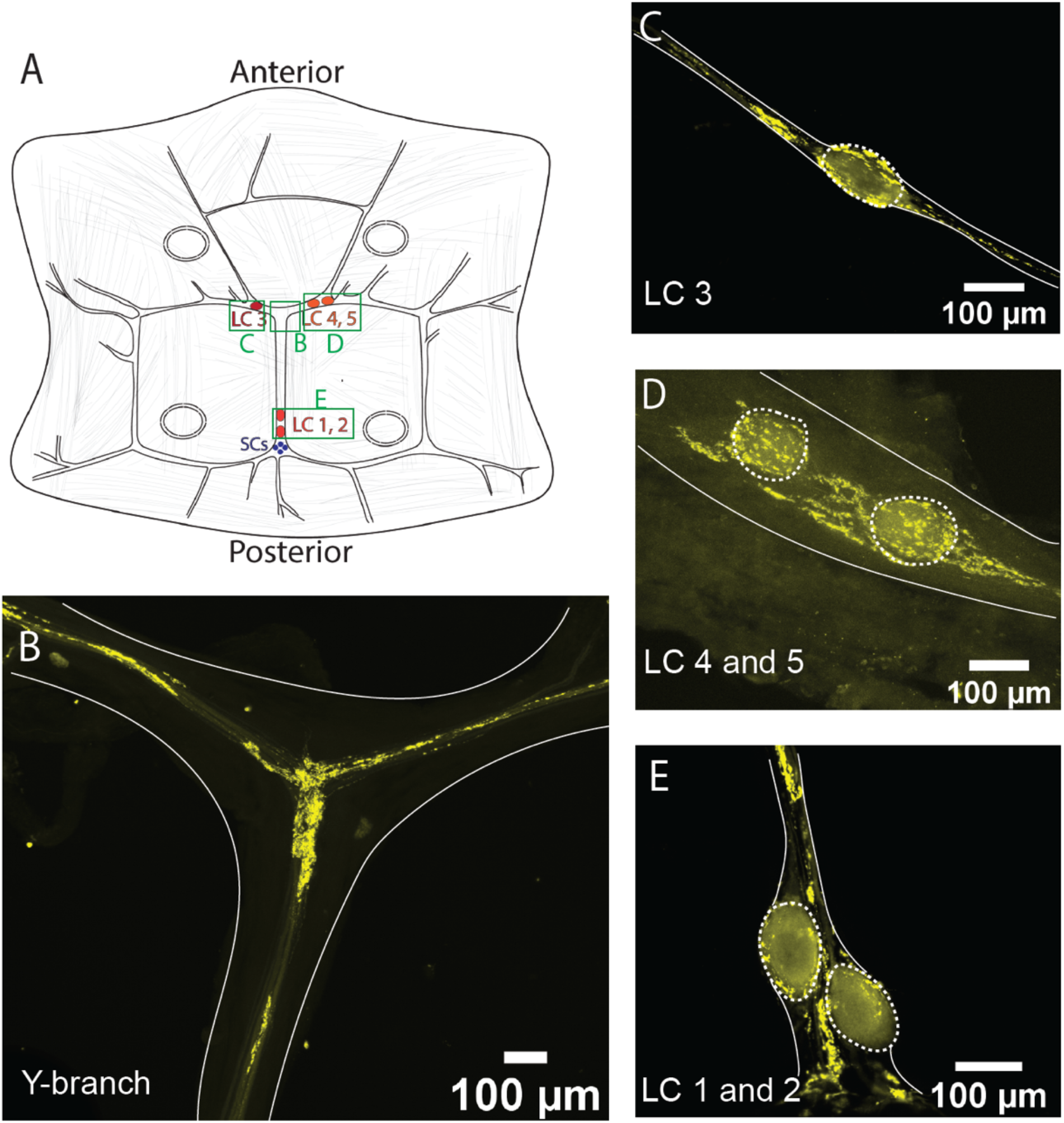
Synapsin-like immunoreactivity (synapsin-LIR) showing the location of pre-synaptic terminals in the CG. Edges of the CG tissue are highlighted in solid white line, and the large cell soma are outlined in dashed ovals. A) Diagram of the CG of *Cancer borealis*, green boxes indicate the areas of interest shown in panels B-E. B) Anterior Y-shaped branch contains a dense neuropil of synapsin-LIR. C) LC3 soma is surrounded by a ring of synapsin-LIR. There are also small clusters of synapsin-LIR to either side of the LC soma. D) LC 4 and 5 display similar clusters of synapsin-LIR around the soma and near the cell bodies. E) Dense synapsin-LIR surrounding LC1 and 2, as well as surrounding the SC and posterior T-shaped branch.

### Proctolin-like immunoreactivity is found throughout the CG

To determine the distribution of different neurotransmitters and the character of the chemical synapses throughout the CG, we performed double immuno-stains for synapsin and a neurotransmitter or neuromodulator of interest.

The immunoreactivity for the excitatory neuropeptide proctolin (proctolin-LIR) is abundant throughout the CG of *C. borealis* (N=8). The proctolin-LIR projections have an intricate branching structure with a beaded appearance (Fig. 3). These projections are found along the trunk and anterior branches of the CG and form dense clusters around the LC bodies. Figure 3A shows a representative pattern of proctolin-LIR surrounding LC3. The innervating proctolin-LIR branches split and form a complex structure surrounding the LC. At the cell body there is a partial overlap between proctolin-LIR and synapsin-LIR. Some synapsin-LIR boutons (yellow) clearly do not contain proctolin-LIR, and some of the proctolin-LIR projections (magenta) surrounding the LC do not overlap with putative chemical synapses. The beaded proctolin-LIR projections are also found at the Y-branch neuropil, and here they also partially overlap with synapsin-LIR (Fig. 3B). The proctolin-LIR projections were clear enough to be traced in most preparations; they extend along the trunk of the CG down to LC1,2 and exit the CG in two bilateral anterior nerves lateral to LC3 and LC4,5 somata.

**Figure 3.**
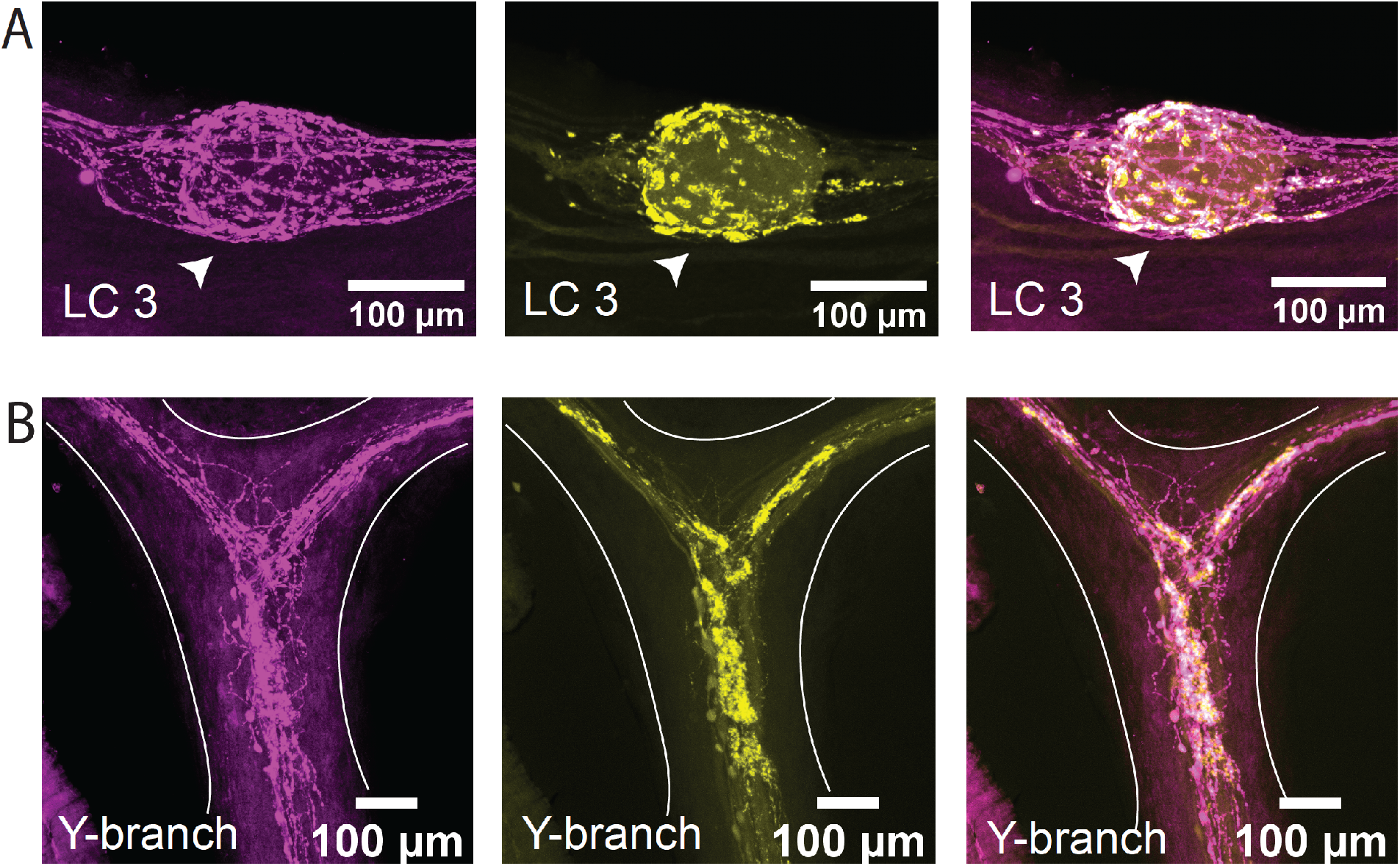
Proctolin-LIR (magenta, left) and synapsin-LIR (yellow, center) in the CG of *Cancer borealis*. The right panel shows the overlay of the two stains, with overlapping regions shown in white. A) Representative LC3 soma (indicated by white arrow) with proctolin-LIR that partially overlaps with synapsin-LIR. Note that not all synapses co-localize with proctolin-LIR signal (yellow synapsin-LIR, right panel), and that not all proctolin-LIR projections overlap with pre-synaptic terminals (magenta proctolin-LIR, right panel). B) Representative Y-branch neuropil where dense proctolin-LIR partially overlaps with synapsin-LIR neuropil.

### GABA-like immunoreactivity is present at nearly all synapses at the LC somata but absent at the Y-neuropil

GABA has a strong inhibitory effect on the rhythm of the CG (Cruz-Bermúdez & Marder, 2007; Kerrison & Freschi, 1992). GABA-like immunoreactivity (GABA-LIR) has been described in the CG of *Panulirus argus* surrounding LC somata (Delgado et al., 2000).

We stained whole-mount CGs with a rabbit polyclonal antibody targeting GABA (N=8). At the large cell somata, the GABA-LIR resembles closely the distribution of synapsin-LIR. Figure 4A shows a representative LC3 with GABA-LIR in magenta. The basket-like structure around the large cell soma is similar to the structure of synapsin-LIR. When the two stains are overlaid, all the synapsin-LIR boutons surrounding the soma also contain GABA-LIR (Fig. 4A, left panel). However, the two stains are not identical. Note the strong GABA-LIR projection under LC3 that extends along the length of the anterior branches of the CG (Fig. 4A, right panel). In Figure 4B one can see the smooth GABA-LIR projections that enter the Y-branch neuropil. Compared with the dense synapsin-LIR at the Y-branch neuropil, there is little GABA-LIR in this area. The overlay of the two stains illustrates how few (if any) of the synapsin-LIR positive boutons at the Y-branch neuropil contain GABA-LIR (Fig. 4B, left panel). The smooth GABA-LIR projections seen in Figure 4 can be easily traced along the nerves of the CG. Similar to the proctolin projections described above, they extend along the trunk of the CG down to LC1,2 and along the anterior branches of the CG until they exit the CG at two bilateral nerves lateral to LC3 and LC4,5.

**Figure 4.**
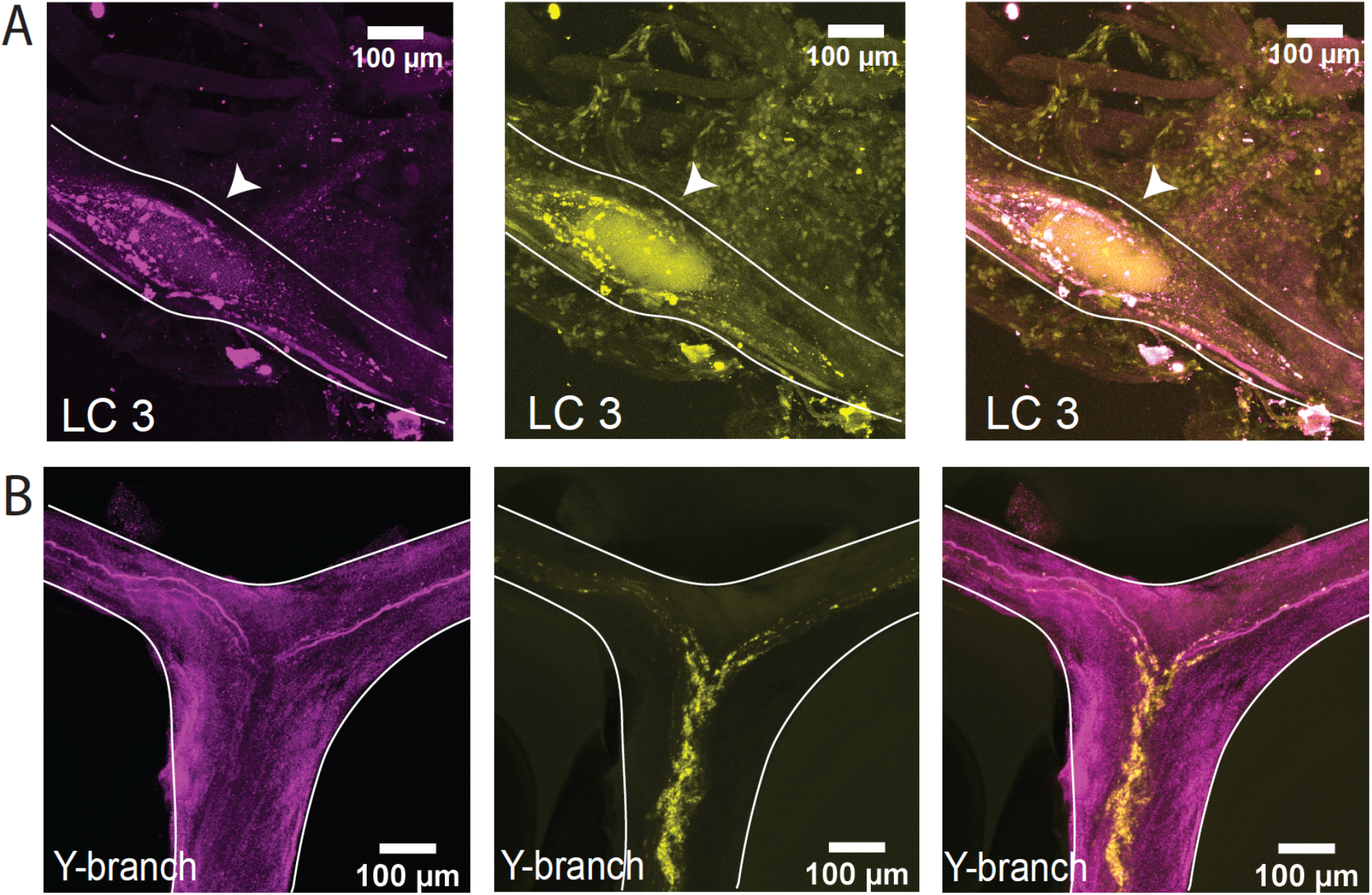
GABA-like immunoreactivity (magenta, left) and synapsin immunoreactivity (yellow, center) in the CG of *Cancer borealis*. The right panel shows the overlay of the two stains, with overlapping regions shown in white A) Representative LC3 soma (indicated by white arrow) where GABA-LIR overlaps strongly with synapsin-LIR, specifically in putative synaptic terminals near the LC soma. Note the strong GABA-LIR projection that passes under the soma of LC3. B) Representative Y-branch neuropil where there is little GABA-LIR signal, except for the salient projections traveling out to the anterior branches (right). Strong immunoreactivity for synapsin is still seen at the Y-branch (center), and there is little overlap between GABA-LIR and synapsin-LIR at the Y neuropil (right).

### Stimulation of nerves containing proctolin-LIR or GABA-LIR affects the CG rhythmic output

Because we found projections exiting the CG for both the excitatory neuropeptide proctolin-LIR and the inhibitory neurotransmitter GABA-LIR, we asked whether the pattern of innervation we observe could be associated with the physiological responses of the CG to stimulation of those nerves. Within the crustacean CG, there are three pairs of innervating nerves which are known to affect the rhythmic bursting of the central pattern generator (Cooke, 2002; Maynard Jr, 1953; Yazawa & Kuwasawa, 1992). These nerves had yet to be formally characterized in *C borealis*. We dissected CGs from the heart muscle while keeping nerves entering the CG intact and pinned the preparation in a Sylgard dish. We used extracellular electrodes to stimulate each nerve in succession and noted the effect on the rhythmic bursting of the CG. Most nerve stimulations had no effect on the bursting behavior of the CG. We stimulated a bilateral pair of nerves on the anterior branches of the trunk lateral to LC3 and LC4,5 (N=5). In the heart muscle, these nerves projected in the posterior direction and join the anterior branches of the CG from the posterior direction. Figure 5A shows a representative extracellular recording from the trunk of a CG while one of these nerves was stimulated: the rhythm frequency increased along with the number of spikes per burst. We fixed and stained a subset of the preparations in which we found an excitatory response for proctolin (N=3) and found clear proctolin-LIR projections within the nerve that excited the CG rhythm (Fig. 5B). This projection connected to the web of proctolin-LIR that encircles the LC soma and innervates the Y-branch neuropil.

**Figure 5.**
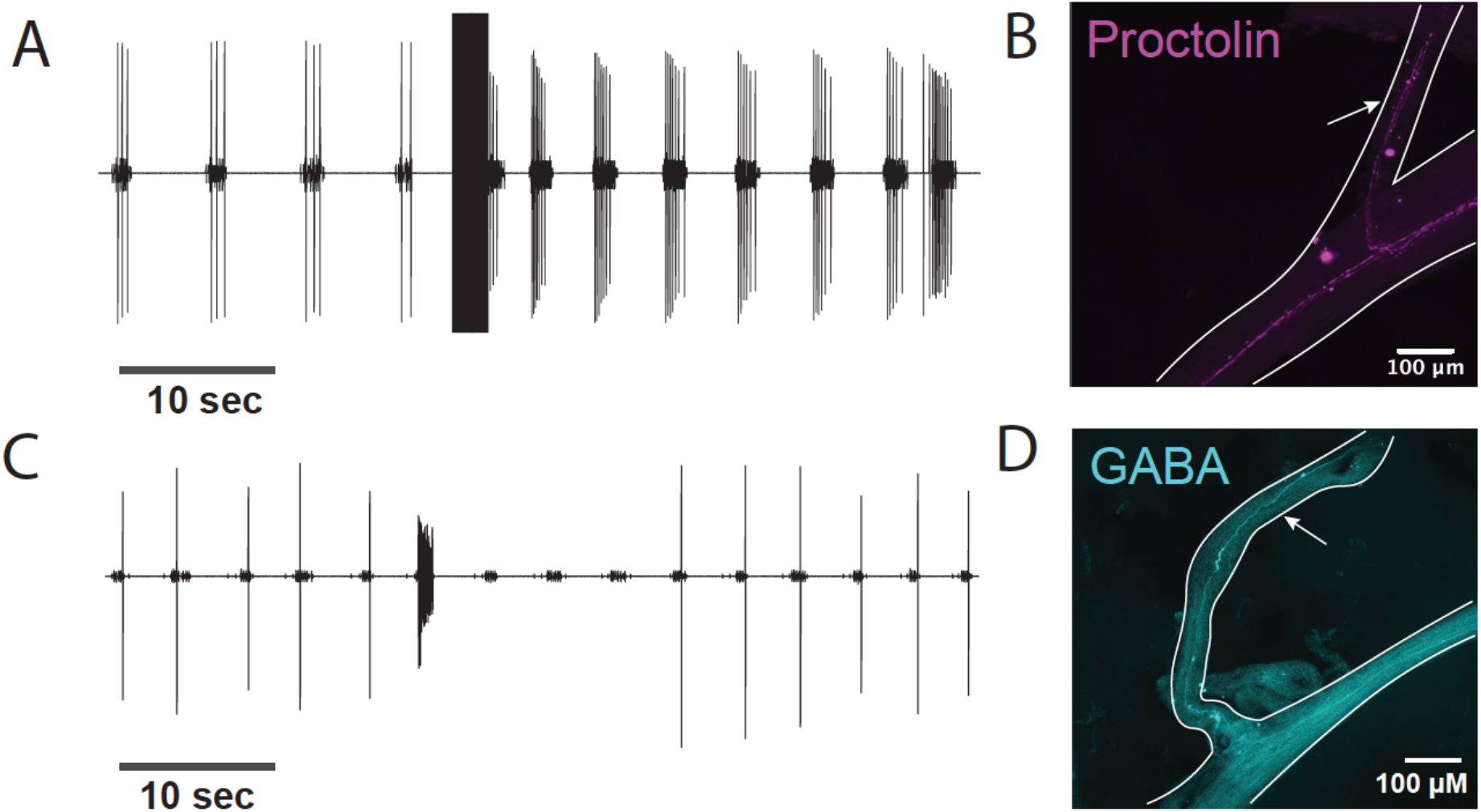
Stimulation of nerves containing Proctolin-LIR or GABA-LIR have respective excitatory and inhibitory effects on CG rhythmic output. (A) Extracellular recording of the rhythmic activity of a CG before and after stimulation of an extrinsic nerve. Stimulation of this nerve resulted in excitation of the CG burst frequency and LC spikes per burst. (B) Proctolin-LIR in the stimulated nerve from the CG recorded from in (A). (C) Extracellular recording of the rhythmic activity of a CG before and after stimulation of an extrinsic nerve. Stimulation of this nerve resulted in inhibition of the CG burst frequency and loss of LC spikes. (D) GABA-LIR in the stimulated nerve from the CG recorded from in (C).

Next, we examined a second pair of nerves that inhibited the CG rhythm when stimulated (N=5). This pair of nerves innervates the CG from the anterior, dorsal side and joins the CG lateral to LC3 and LC4,5. Figure 5C shows a representative recording from the CG when one of these nerves was stimulated: the bursting of the LC stopped for several cycles, and the rhythmic bursting of the SC was slowed. We fixed and stained a subset of the CG in which we found an inhibitory response using anti-GABA primary antibodies (N=3). Clear GABA-LIR projections exited the CG at the inhibitory nerves (Fig. 5D). These smooth GABA-LIR projections connected to the projection that extends along the branches of the CG and surrounds the LC somata.

### Variable distribution patterns and overlap with putative chemical synapses for other neuromodulators in the CG

The neuromodulator allatostatin-B (AST-B) is a neuropeptide found in the CG (DeLaney & Li, 2020). Allatostatin-B-like immunoreactivity (ASTB-LIR) within the CG of *C. borealis* was most prevalent at the LC soma and the Y-branch neuropil, similar to the distribution of synapsin-LIR (Fig. 6). Note that at the LC soma the ASTB-LIR pattern closely overlaps with the synapsin-LIR (Fig. 6A, representative cell bodies of LC4 and LC5). ASTB-LIR around the cell bodies is punctate, and with no clear projections along the branches of the CG. This punctate structure is also present at the Y-branch neuropil, where some putative synapses, but not all, overlap with the ASTB-LIR (Fig. 6B).

**Figure 6.**
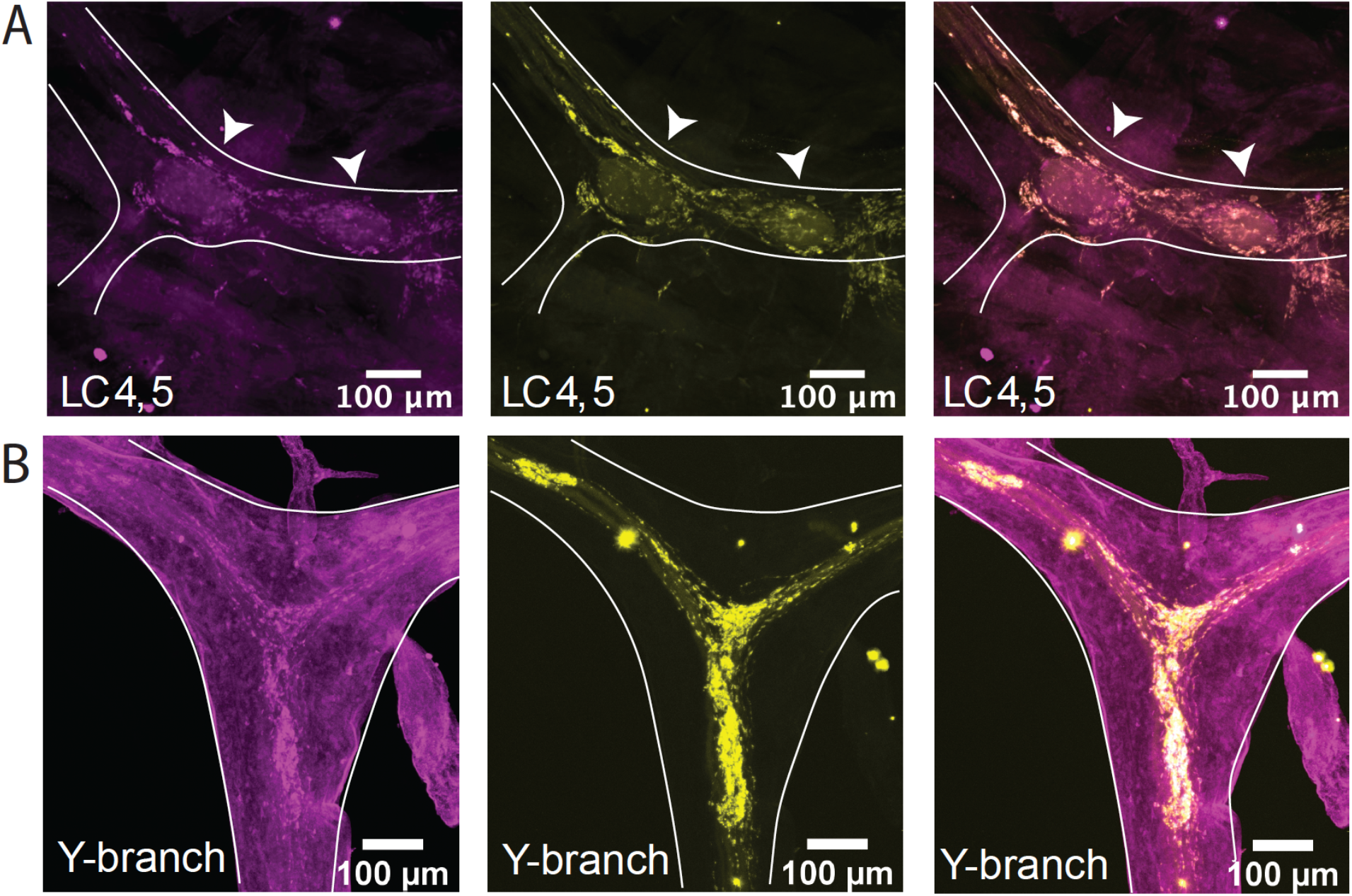
AST-B-like immunoreactivity (magenta, left) and synapsin immunoreactivity (yellow, center) in the CG of *Cancer borealis*. The right panel shows the overlay of the two stains, with overlapping regions shown in white A) Representative LC4 and 5 somata (indicated by white arrows) where ASTB-LIR overlaps strongly with synapsin-LIR, specifically in synaptic terminals near the LC soma. B) Representative Y-branch neuropil where the ASTB-LIR signal partially overlaps with the synapsin-LIR at the Y-branch.

We also found immunoreactivity for excitatory neuromodulators red pigment concentrating hormone (RPCH, N=5) and FLRFamide peptides (N=9) within the CG. Unlike the extensive proctolin-LIR, these neuromodulators have a much more punctate and diffuse pattern of immunoreactivity (Fig. 7). More than 30 biologically active FLRFamides have been identified in *C. borealis* (Marder et al., 1987; Trimmer et al., 1987; Weimann et al., 1993).The antibodies used here do not distinguish among many extended FMRFamdie and FLRFamide peptides Mass spectrometry imaging found many FLRFamides are present in the CG, and it is likely that many are biologically active (DeLaney & Li, 2020). Small puncta of FLRFamide-like immunoreactivity (FLRF-LIR) can be seen around the LC bodies (representative LC3, Fig. 7A) and at the Y-branch neuropil (Fig. 7B). All puncta of FLRF-LIR overlap only partially with the synapsin-LIR within the same CG.

**Figure 7.**
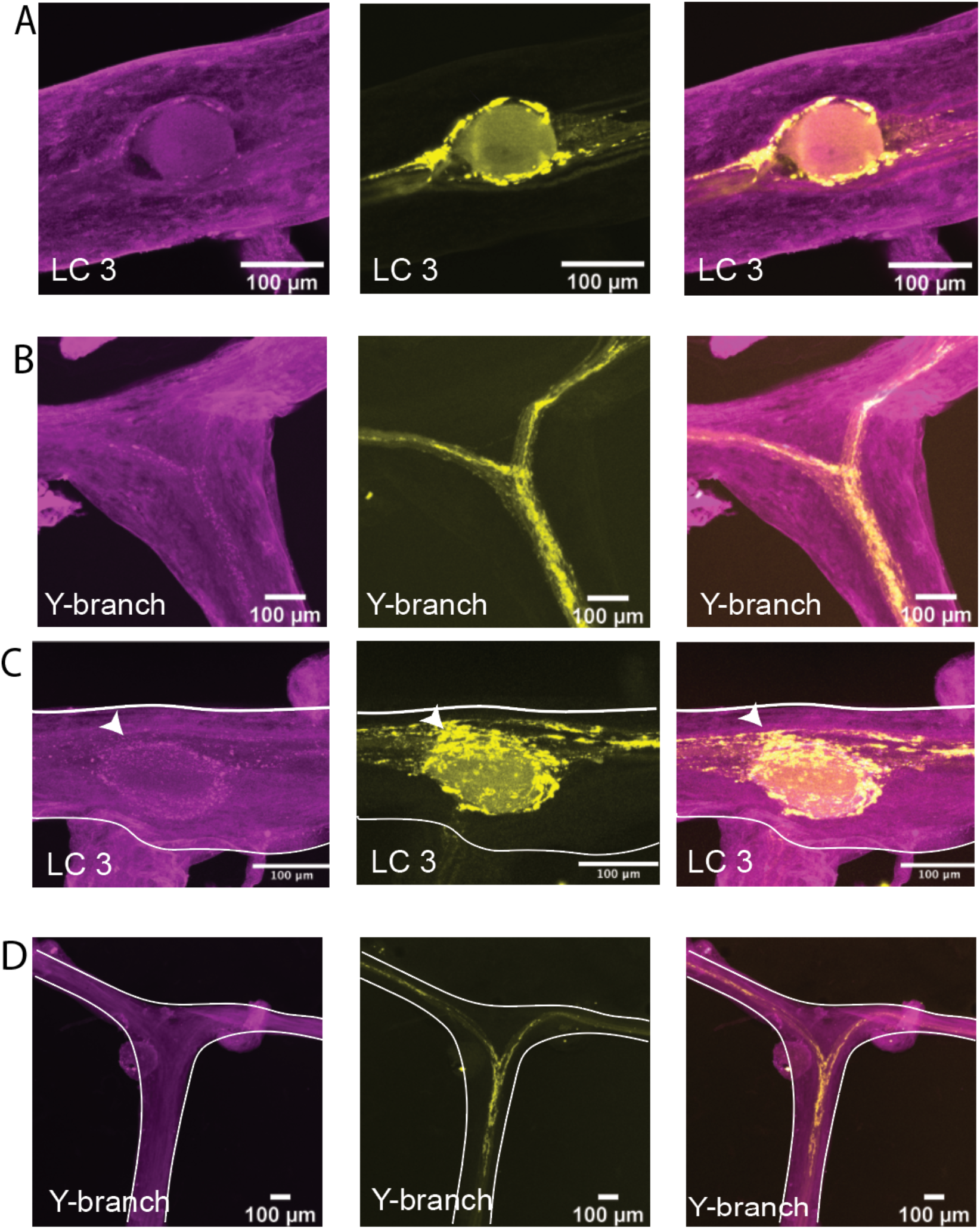
FRMFamide-like immunoreactivity (A-B, magenta, left), RPCH-like immunoreactivity (C-D, magenta, left) and synapsin immunoreactivity (yellow, center) in the CG of *Cancer borealis*. Both excitatory neuromodulators have similar patterns of immunoreactivity and overlap with synapsin-LIR in the CG. Both neuropeptides have weak immunoreactivity surrounding the LC somata and partially overlap with synapsin-LIR around. The soma of LC3 (A and C). FMRFamide-i is present at the Y-branch neuropil and again partially overlaps with synapsin-LIR there (B). RPCH-LIR is not found at the Y-branch neuropil (D).

The neuropeptide RPCH (pQLNFSPGW-NH_2_) is found throughout the nervous system and endocrine system of crustaceans (Dickinson et al., 2001; Dickinson & Marder, 1989; Dickinson et al., 1993). It is excitatory within the crustacean CG and activates similar excitatory currents to the neuropeptide proctolin (Cruz-Bermúdez & Marder, 2007; Swensen & Marder, 2000). RPCH-like immunoreactivity (RPCH-LIR) is found surrounding the LC soma in distinct punctate structures (Fig. 7C) that overlap partially with synapsin-LIR near the soma. Although we found RPCH-LIR near the LC neuropils, there was no RPCH-LIR at the Y-branch neuropil (Fig. 7D).

Overall, each neuromodulator we stained for showed different patterns of immunoreactivity throughout the CG. Each had only a partial overlap with putative chemical synapses around the LC somata. While proctolin-LIR and FLRFamide-LIR were abundant at the Y-branch neuropil, RPCH-LIR and GABA-LIR are absent there, indicating that there may be a physical separation of the site of action for different neuromodulators within the CG of *Cancer borealis*.

## Discussion

All neuronal circuits are affected by neuromodulation. Within a network, neuromodulators can act individually, as co-transmitters, and/or hormonally through circulation (Marder, 2012). Work on small circuits in which neuromodulation is well defined illustrates that the location and means of release of neuromodulators is important and can determine their effect on the circuit (Blitz et al., 1999; Nusbaum et al., 2017; Nusbaum et al., 2001). In this study we set out to characterize the putative location of chemical synapses within the CG and describe the distribution of some of the neuromodulators that may act at those synapses. The distribution of proctolin-, RPCH-, FLRFamide- and CbAST-B1-like immunoreactivity within the crustacean CG are described here for the first time. In addition, we observed the pattern of GABA immunoreactivity within the CG of *C. borealis*. While we examined only a subset of the neuromodulators known to act on the CG, their differences in distribution suggest specialized regions of integration for different signaling molecules.

### Sites of chemical synaptic signaling in the CG

Among invertebrates, the majority of synapses are found between axons and dendrites, and early studies posited that crustacean cell bodies receive no direct synaptic input, and that all integration occurs within the dendritic arbor (Evoy, 1977). Therefore, the clustering of putative chemical synapses around the somata of the large motor neurons in the CG is an unusual configuration. However, this observation is consistent with previous electrophysiological and morphological studies in the CG.

The specifics of pattern generation in the CG may affect the efficacy and practicality of somatic synapses. The somata and proximal axon of CG motor neurons are not capable of initiating action potentials and are thought to be the site of extended integration of dendritic impulses (Benson & Cooke, 1984). The motor neurons of the CG are very large (∼100µm in diameter), and as such the cell body of the neuron provides a large surface for synapses to form. Crustacean motor neurons can be electrotonically compact, and synaptic potentials up to 800µm away from the recording sites have been shown to propagate faithfully (Otopalik et al., 2017). Therefore, for these neurons the exact location of synaptic inputs may be less important for global summation at the site of action potential initiation. The CG is also embedded in the heart muscle and stretch sensitive dendrites extend from the cell body and proximal axon that provide excitatory input when the muscle is stretched (Alexandrowicz, 1932, 1934; Hartline, 1967; Sakurai & Wilkens, 2003). This organization could enhance phasic input to the CG. Finally, each neuron within the CG has an intrinsic driver potential that allows for spontaneous rhythmic bursting activity (Cooke, 2002; Wantabe, 1958). Taken together, the summation of the driver potential with stretch feedback distal to the impulse generation zone implies that inhibitory or excitatory somatic inputs could have a strong effect on the overall output of the CG (Cooke, 2002). In electron microscopy studies of the crab *P. sanguinolentus* CG, Mirolli et al. (1987) noted synapses on short projections that may have extended directly from somata of the large motor neurons. These small projections from the LC soma could also serve to increase the effectiveness of any somatic synaptic inputs (Mirolli et al., 1972).

In his early studies of the anatomy of crustacean CG, Alexandrowicz (1932) noted that the dendrites of both the large motor neurons and small premotor neurons branched extensively at the anterior branch of the CG trunk. Combined with these anatomical descriptions, the dense synapsin-LIR neuropil we found at the Y-branch is likely to be a site of extrinsic modulation for both the LC and SC. This is consistent with the hypothesis that extrinsic modulation is also integrated in the proximal axon of the motor neurons and modulates the driver potential of the LC prior to impulses reaching the trigger zone for action potentials (Benson & Cooke, 1984).

### Extrinsic modulation of the CG

At least three pairs of extrinsic nerves innervate the CG and modulate its rhythmic output. Initial studies of the CG confirmed that one of these nerves is inhibitory and may contain the neurotransmitter GABA (Alexandrowicz, 1934; Delgado et al., 2000; Maynard, 1961). The identity of neuromodulators in the CG’s excitatory nerves is relatively unexplored. In this study, we confirmed the presence and localization of at least two pairs of modulatory nerves to the CG of *C. borealis* through a combination of electrical stimulation and immunohistochemistry. By combining these two techniques, we confirm that GABA-LIR is present on nerves that inhibit the output of the CG when stimulated, as had been postulated in studies of other crustacean species. Furthermore, we also identified proctolin-LIR within nerves that excited the CG rhythm when stimulated. This study therefore provides evidence that proctolin is among the neuromodulators released locally through the excitatory nerve. However, further physiological study is necessary to determine whether proctolin release from this nerve is necessary and sufficient to excite the CG rhythmic output. There are many other candidates for excitatory neuropeptides that could be carried on this nerve and co-released with proctolin that were not stained for here. In particular, staining for crustacean cardioactive peptide (CCAP) in the central nervous system of *C. sapidus* implicates CCAP as another excitatory neuromodulator that may signal through this nerve (Fort et al., 2007).

### Neuromodulation and possible co-modulation at putative somatic synapses

Most of the neuromodulators we stained for were localized to the putative somatic synapses on the LC. As noted above, the GABA-LIR and ASTB-LIR were particularly dense at the putative somatic synaptic sites. The localization of GABA-LIR in the CG of *C. borealis* supports the idea that GABA input is specifically localized near large cell somata for effective inhibition. B-type allatostatins are part of a major neuropeptide family that is widely conserved across decapods. Many isoforms for ASTs exist both across species and within species. At least eight types of AST-B are found in *C. borealis* (DeLaney & Li, 2020; Fu et al., 2005; Fu & Li, 2005). Because of conserved domains in the CbAST-B family, the ASTB-LIR we observed could combine signals from many different peptide isoforms present in the CG of *C. borealis* (DeLaney & Li, 2020). Careful MALDI imaging would be required to confirm any localization differences between these peptide types. The actions of AST-B on the crustacean pyloric rhythm are known to be state dependent and vary widely among individuals (Fu et al., 2007), leading to the speculation that AST-B could be an important co-modulator that acts in combination with other neuroactive compounds. Given that there is significant overlap between GABA-LIR and synapsin-LIR at LC soma synapses, and ASTB-LIR and synapsin-LIR at the LC somata, it is likely that there is co-localization of GABA and AST-B. These results support the theory that CbAST-B1 may be a neuromodulator whose effects are only measurable when co-released (Michael P Nusbaum et al., 2001).

Immunoreactivity for the excitatory peptides proctolin and RPCH were also found at the putative LC somatic synapses. Proctolin is known to differentially affect the SC and the LC in the CG, which is consistent with our finding that proctolin-LIR is found surrounding both the LC somata, SC somata and at the Y-branch neuropil (Saver et al., 1999; Sullivan & Miller, 1984). The additional presence of RPCH-LIR at putative LC somatic synapses indicates that there may also be co-modulatory release of excitatory neuropeptides from these sites.

### Multi-modulated pattern generation

The presence of so many neuromodulators within the CG suggests that it is an important target for physiological regulation, and we know that the output of the CG *in vivo* can be highly variable depending on environmental context (Kushinsky et al., 2019). It is easy to imagine why cardiac output would need to be highly flexible depending on the needs of the animal, which may be one reason why there are so many neuromodulators that act on the cardiac nervous system (Cruz-Bermúdez & Marder, 2007). Here we illustrate another way in which modulation can be targeted, where some neuromodulators are preferentially localized to somatic synapses. This segregation of inputs will likely affect how and in what contexts a neuromodulator will affect the output of the CPG, enhancing the flexibility of neuromodulation within the circuit.

## Acknowledgments

Supported by NIH R35 NS097343 and F31-NS113383.

